# Differential carbohydrate utilization and organic acid production by honey bee symbionts

**DOI:** 10.1101/294249

**Authors:** Fredrick J. Lee, Kayla I. Miller, James B. McKinlay, Irene L. G. Newton

## Abstract

The honey bee worker gut is host to a community of bacteria that primarily comprises 8-10 bacterial species. Collectively, these microbes break down and ferment saccharides present in the host’s diet. The model of metabolism for these gut symbionts is rooted in previous analyses of genomes, metagenomes, and metatranscriptomes of this environment. Importantly, there is a correlation between the composition of the gut microbiome and weight gain in the honey bee, suggesting that bacterial production of organic acids might contribute to the observed phenomenon. Here we identify potential metabolic contributions of symbionts within the honey bee gut. We show significant variation in the metabolic capabilities of these microbes, highlighting the fact that although the microbiota appears simple and consistent based on 16S rRNA gene profiling, strains are highly variable in their ability to use specific carbohydrates and produce organic acids. Finally, we confirm that the honey bee core microbes, especially a clade of γ-proteobacteria (i.e. *Gilliamella*), are highly active *in vivo*, expressing key enzymatic genes critical for utilizing plant-derived molecules and producing organic acids. These results suggest that *Gilliamella*, and other core taxa, may contribute significantly to weight gain in the honey bee, specifically through the production of organic acids.

## Introduction

Insects are some of the most abundant and diverse species on earth, inhabiting a plethora of unique environments, ranging from the arid sands of the hottest desert to frigid temperatures of glaciers (1). In order to thrive in such unusual environments, many species of insects have evolved specialized symbioses with microbes capable of synthesizing essential nutrients (*e.g.*, amino acids and vitamins) or degrading complex macromolecules (2). The world’s most important agricultural pollinator, the honey bee (*Apis mellifera*), is no exception, as it relies on microbes to aid in the digestion of its plant-derived diet of honey and bee bread (3). After consumption, food ingested by the bee is metabolized by the microbes that inhabit the honey bee gut, which include a small number of dominant taxa that constitute bacterial species considered to be “core” to the honey bee gut microbiota, including the proteobacteria *Snodograssella alvi* (Betaproteobacteria), *Parasaccharibacter apium* (Alphaproteobacteria)*, Frischella perrara* and *Gilliamella apicola* (Gammaproteobacteria), two *Lactobacillus* species (Firmicutes), a *Bifidobacterium* species (Actinobacteria), and a rarer *Bacteroidetes* (4). Three of these phyla, the Gammaproteobacteria, Firmicutes, and Bifidobacteria, dominate predicted functional capacity of the honey bee gut microbiome, based on a metatranscriptomic survey on the honey bee gut (5). The current proposed model of bacterial community metabolism in the bee gut, based on annotated gene presence and expression, suggests that these core microbes participate in the breakdown and fermentation of host dietary macromolecules into an array of alcohols, gases, short chain fatty acids (SCFAs), and other organic acids, some of which likely serve to sustain the host (5–10). Complementing this work, some studies have demonstrated, *in vitro*, the ability of some strains to ferment a variety of substrates (5, 9, 11–14) and produce organic acids (i.e. lactate and acetate)(11, 14). In addition to these observations, a separate study also recently demonstrated that gnotobiotic, mono-colonized bees possessed guts containing significant depletion in plant derivatives (i.e. flavonoids, ω-hydroxy acids, and phenolamides), likely originating from pollen consumed by the host, and an enrichment in fermentative products in the form of organic acids and SCFAs, compared to bees with a depleted microbiota (15). The production of these fatty acids may contribute to honey bee health. Indeed, a recent study identified a correlation between host weight gain and organic acids in the digestive tract of bees (3). Bees supplemented with gut bacteria showed an increase in the variety and amount of organic acids present within the host digestive system, as well as a significant increase in host weight gain and the expression of host genes related to insect growth and nutrient homeostasis, compared to controls. Collectively, these results link the honey bee symbionts to the production of organic acids and then to host growth and nutrient acquisition. It has been proposed that honey bee core bacteria likely convert carbohydrates in the bee diet to a variety of organic acids, which can either act as a signaling molecules eliciting a host response, or act as building blocks for host derived anabolism.

In light of these recent findings, we sought to determine the metabolic contribution of the Gammaproteobacteria, Firmicutes, and Bifidobacteria, within the bee gut, as these phyla dominate the metatranscriptomic profile of the digestive tract of the honey bee. Although there are well-defined species within these groupings, there is still a large diversity of honey bee strains found within individual bees and colonies (16). The strain diversity within each of the honey bee core clades is reflected in differences in genomic content (7, 16), suggesting that significant functional diversity may exist within each of the core 8-10 clades associated with the honey bee. In this study, we address fundamental questions regarding metabolic characteristics of the honey bee gut microbiota, such as: 1) What is the pattern of anaerobic utilization of environmentally relevant carbon sources across dominant symbionts, and what are the direct byproducts produced, per taxa, *in vitro*? 2) Are there functional differences in the patterns of utilization between isolates of the same species? 3) What contribution does an individual taxon make to the utilization and the subsequent fermentation of environmentally relevant substrate throughout the digestive tract of the honey bee? Through utilizing a combination of molecular, biochemical, and physiological approaches, both *in vitro* and *in vivo*, we perform an in-depth metabolic analysis on 17 isolates from the honey bee gut. Our results suggest that overall, bee-associated microbes are primed to metabolize certain components of the honey bee diet, but that significant variation exists within clades, suggesting that each member contributes differentially to honey bee metabolism.

## Materials and Methods

### Bee Sampling and Bacterial Culture

Worker honey bees were collected from colonies located at an apiary in Bloomington, IN and transported to lab in sterile vials on ice. Hindgut dissections were performed using aseptic technique and sterile equipment. Whole digestive tracts of 2-3 bees were homogenized in 1X phosphate buffered saline (PBS) with a sterile disposable pestle and a dilution series was plated onto Brain-Heart Infusion (BHI), de Man, Rogosa and Sharpe (MRS), and Luria-Bertani (LB) agar (BD Difco™). All agar plates were incubated under anaerobic conditions and supplemented with CO_2_ using the GasPak EZ system (BD, New Jersey) at 37°C for 4 – 5 days. Isolated colonies were subcultured on agar before genotyping (below).

### 16S rRNA genotyping of bacterial isolates

For each isolate, genomic DNA was extracted using the QIAGEN DNeasy Blood & Tissue kit with one modification: the inclusion of a bead-beating step before lysates were loaded onto the Qiagen column. DNA was quantified using a spectrophotometer (BioTek instruments) and used in PCR using Phusion^®^ High Fidelity PCR mix with HF buffer (New England BioLabs INC.), with 27F (5’-AGA-GTT-TGA-TCC-TGG-CTC-AG-3’) and 1492R (5’–ACG-GCT-ACC-TTG-TTA-CGA-CTT-3’) primers. A reaction consisted of 0.4 mM primers, 1x Phusion mix, PCR grade water, and 1 µL of sample DNA, under the following conditions: 5 min at 98°C; 35 cycles-10 sec at 98°C, 30 sec at 55°C, 30 sec 72°C; Extension-10 min at 72°C. PCR products were visualized using gel electrophoresis and all isolates with PCR products of the correct size were used for sequencing (Beckman Coulter Genomics). Resulting ab1 files were concatenated and exported to Fasta. To align, trim, and classify sequences, Mothur v.1.33.3 software was used with the following commands (align.seqs, trim.seqs, and classify.seqs), using a custom reference database (Greengeen + honey bee specific dataset) (Newton and Roeselers, 2011) for classification. To demonstrate the evolutionary relationship between isolates, a 16S rRNA gene phylogeny was constructed, using the concatenated 16S rRNA gene sequence used for classification. The tree was rooted with *Aquifex aeolicus* (Accession: AJ309733.1) and to serve as representatives of the four dominant bacterial families within the honey bee (*Gilliamella apicola* - Accession: NR_118433.1 (Gilliamella), *Lactobacillus* sp. wkB8 - Accession: NZ_CP009531 - Locus Tag: LACWKB8_RS00480 (Firm-5), Uncultured *Lactobacillus* sp. - Accession: HM113352.1 (Firm-4), and *Bifidobacterium asteroides* strain Hma3 - Accession: EF187236.1 (Bifido) were used. Sequences were uploaded to SINA 1.2.11 aligner (ARB SILVA), and aligned using the default settings, after which the FASTA formatted alignment was converted to MEGA format (MEGA v6.06) and a maximum likelihood tree (GTR + G + I) was constructed using 1000 bootstrap replicates. Divergence estimates were performed in BLASTClust (Alva et al, 2016).

### Carbohydrate Utilization & Byproduct Production of Isolates

To prepare cell suspensions to assay substrate utilization and product formation, all isolates classified as actinobacteria or bacilli were grown on MRS agar, while gamma-proteobacteria isolates were grown on BHI; these media correspond to the medium under which the microbes were isolated. Broth cultures were grown anaerobically at 30°C for 6 days, after which cultures were pelleted via centrifugation and the supernatant removed. The pellet was washed with salt solution (137.0 mM NaCl, 2.7 mM KCl, pH 6.0) to remove remaining media. Bacilli and actinobacteria, grown in MRS, were washed twice to decrease the transfer of nutrients present in the spent MRS media. Isolates were normalized for growth based on OD, across taxonomic classes, such that isolates with more robust growth were diluted in a larger amount of salt solution.

To test the ability of isolates to utilize carbon substrates, we used the MT2 MicroPlate™ assay. In the assay, each well in a 96-well plate contained an aliquot of a TTC dye. To each well, 150 µL of cell suspension from each isolate was added, with or without 15 µL of a filter-sterilized carbon source (either 1.1 M of D-(+)-glucose (SIGMA G8270), D-(-)-fructose (ACROS Organics 57-48-7), D-sucrose (Fischer Scintific BP220-1), D-(+)-mannose (SIGMA M6020), L-rhamnose (SIGMA W373011), D-(+)-galactose (SIGMA G0625), D-(+)-xylose (SIGMA X1500); or 13.1% D-(+)-cellobiose (SIGMA 22150), or 2.1% pectin (SIGMA – Pectin from apple – SIGMA 76282) solutions). Negative controls received 15 µL of salt solution. To verify that substrate solutions were void of microbes and that cross-contamination between wells was not occurring, an additional negative control was filled with 15 µL of a sole carbon source with 150 µL of salt solution. Each experimental condition and control was analyzed in triplicate within a single plate. 96 well plates were covered with an optical adhesive film, and incubated anaerobically at 30°C for 3 days. To measure reduction of TTC (and utilization of a substrate), absorbance readings at 590 nm were taken, in triplicate, for each well. Significant utilization was determined by performing a t-Test comparing 590 nm readings between experimental (cell suspension + 15 µL substrate) and negative control (cell suspension + 15 µL of salt solution) wells. All statistics were performed in IBM SPSS Statistics Version 24 on datasets tested for a normal distribution. Biological triplicates for each condition or control were then combined in a screw-top cryovial and stored at -80°C until high performance liquid chromatography (HPLC) analysis (below).

To determine the soluble fermentation products, pooled samples were thawed, vortexed, and then centrifuged at 13,000 RPM for 5 minutes to pellet bacterial debris. After centrifugation, samples were filtered through a 0.45 µm HAWP membrane (Millipore Corp., Bedford, Mass.) and acetate, lactate, formate, and succinate were quantified by using a Shimadzu HPLC (Kyoto, Japan) as previously described (17).

### Assaying the expression of enzymatic genes

For each dissection, the crop, midgut, and hindgut were aseptically incised from the full digestive tract, and placed in a 2.0 mL Lysing matrix E tube with 500 uL of Trizol^®^ reagent. Samples were homogenized using a FastPrep^®^-24 instrument, twice, at 6.0 m/s for 40 seconds. RNA were extracted from samples using the standard protocol provided by the manufacturer. After the final washing, RNA was resuspended in RNA Storage Solution (Bioline) and stored at -80°C for downstream analysis. For each gut chamber, each biological replicate consisted of three separate individuals. All analyses were performed in triplicate with the same biological samples.

Primer sets specific for a particular enzymatic gene and group of honey bee microbes (Gilliamella, Firm-5, and Bifido), were designed for qRT-PCR analyses (Supplementary Table 1, Supplementary Figure 2). Selected primer sets fulfilled the following requirements: 1) they amplified a product of the expected size, 2) the resulting sequenced product had a top BLAST hit(s) to the intended bacterial taxa and the gene of interest, or for the reference sequence used to design the primer set and 3) they showed specificity for the bacterial group of interest in laboratory experiments (Supplemental Table 1, Supplemental Figures 2). RNA extracted from individual gut chambers (foregut, midgut, and hindgut) were used as a template for qRT-PCR reactions. Prior to running reactions, all RNA samples were treated with RNase-free DNase I (Ambion) as prescribed by the manufacture. RT-qPCR was performed using SensiMix ™ SYBER R Hi-ROX One-Step kit and protocol. As a reference locus in these analyses, we used the 16S rRNA gene, using a universal bacterial primer set (S-*-Univ-05150a0S-19/S-D-Bact-0787-b-A-20 (18)). Reactions were run on an Applied Biosystems StepOne RT-PCR system: Initial denaturation-5 min at 98°C; 40 cycles -15 sec at 98°C, 1 min at 60°C, 30 sec 72°C; Extension-10 min at 72°C. A standard curve was generated for each primer set separately, to account for primer efficiencies in our analyses. All statistics were performed using IBM SPSS Statistics Version 24.

**Table 1.** Cluster analysis of the 16S rRNA gene sequences of 17 cultured honey bee gut isolates used for metabolic analysis. A FASTA file with near full-length 16S rRNA sequences for each isolate was uploaded to BLASTClust (Alva et al., 2016). Standard settings were used for the analysis, using 97% or 98% identity thresholds. The total number of isolates analyzed for each class of bacteria is provided, along with the number of phylotype clusters at 97% and 98% ID.

| Bacterial Class | Total Number of isolates | Number of Phylotypes (%ID) |  |
| --- | --- | --- | --- |
|  |  | 97 | 98 |
| **γ-Proteobacteria** | 5 | 5 | 5 |
| **Bacilli** | 6 | 5 | 6 |
| **Actinobacteria** | 6 | 1 | 6 |

To quantify the expression of metabolic genes from honey bee specific clades of bacteria, across gut chambers of the honey bee, Ct values acquired from the quantification analyses of individual genes were initially normalized to the mean amplification of the reference locus (16S rRNA gene) across gut sections. The advantage of normalizing the data to 16S RNA gene is that it demonstrates the presence of genes, relative to a proxy for the whole bacterial community present throughout the digestive tract of the honey bee. Next, for each metabolic gene prescribed to a taxa of interest, we standardized the data to the gut section with the lowest observed amplification (i.e the crop) for calculating the relative expression (RNA). To determine the relative ratio of expression of metabolic loci attributed to a particular taxa, across gut sections, normalized datasets were standardized to the gut section and locus with the lowest level of expression. To determine variation in the relative abundance, relative expression, or relative ratio of expression of loci across gut sections, we performed an ANOVA or Kruskal-Wallis test on the standardized data acquired for each individual gene analyzed, which was dependent on results obtained from normality test of individual gene datasets. Results with statistical significance (p ≤ 0.05) were subjected to pairwise comparisons using one-way ANOVA/post hoc or Mann-Whitney test.

## Results

### The honey bee core microbiota utilizes carbohydrates from the host diet

To determine the ability of gut isolates to utilize environmentally relevant substrates, we isolated 17 honey bee associated bacteria, spanning three transcriptionally dominant clades: *Gilliamella*, *Lactobacillus* Firm-5, and *Bifidobacterium* (Figure 1, Supplemental Figure 1). Based on the construction of a 16S rRNA gene phylogeny, including data from the 17 cultured isolates and 145 sequences previously obtained from bee-associated microbiota, some of the microbial cultures used herein are closely related to core honey bee bacteria while others fall outside of the core clades (Supplemental Figure 1). When comparing the diversity within the clades (based on 16S rRNA genes), all isolates are unique at 98% and 99% divergence, and some are unique at 97% divergence (Table 1), indicating that these microbes represent different bacterial strains and species.

**Figure 1.**
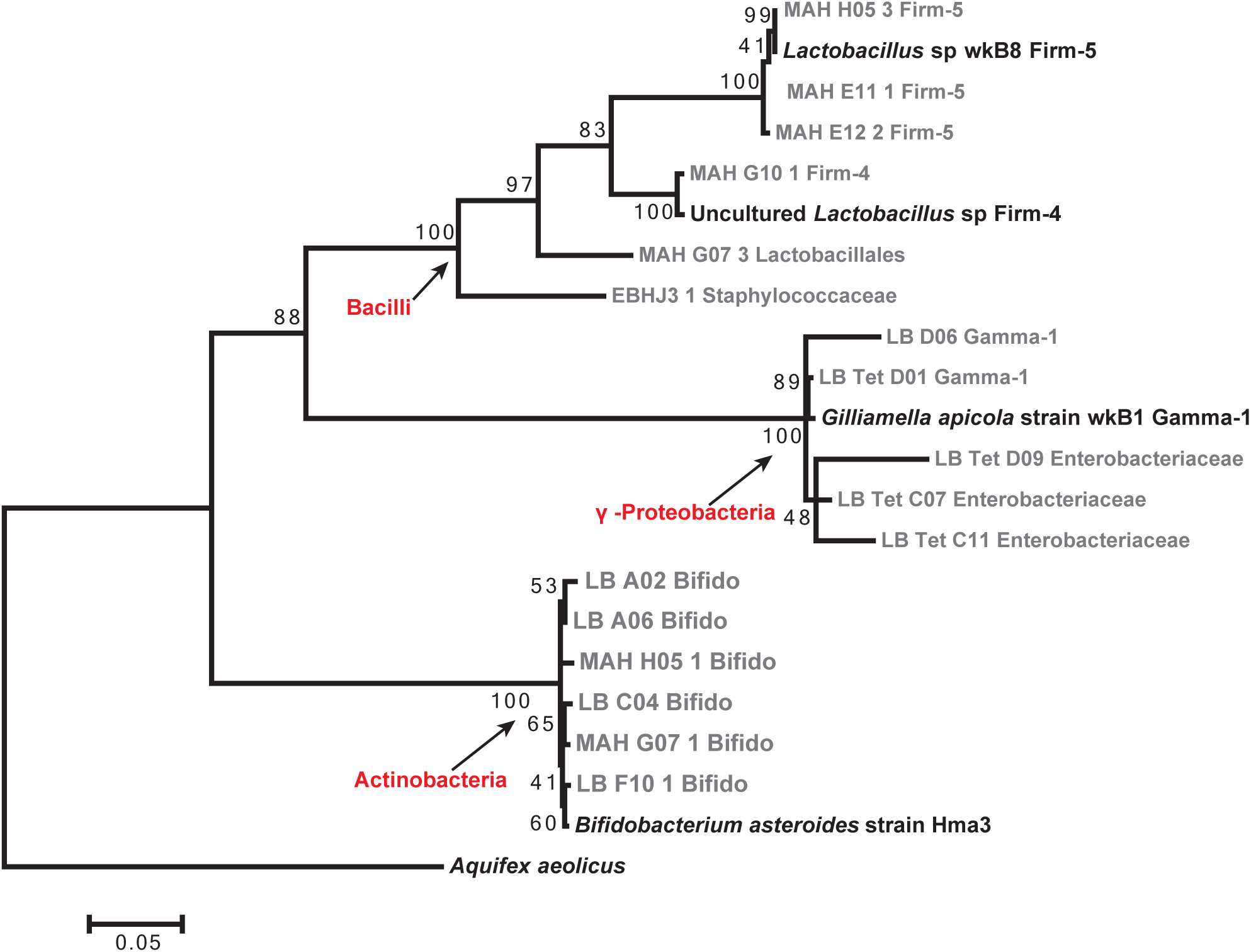
16S rRNA gene phylogeny of gut isolates selected for metabolic analyses, in relationship to reference sequences of honey bee specific taxa. Sequences were aligned using SINA Aligner 1.2.11, the output file was used to construct a Maximum likelihood phylogeny, in MEGA 6.06 with GTR (G+I) and 1000 bootstraps. Bootstrap support is provided at nodes. Reference sequences indicated by black text. Cultured isolates indicated by grey text. Taxonomic class highlighted by arrow.

To test the ability of gut bacteria to utilize sole carbon sources relevant to the honey bee diet, we used Biolog MT2™ plates, containing an aliquot of an indicator tetrazolium chloride dye (TTC), that when combined with a sole carbon source, along with an aliquot of culture, detects substrate utilization via a concomitant color change due to reduction of the dye. Each isolate was examined for the sole utilization of 9 carbon sources (glucose, fructose, sucrose, xylose, pectin, cellobiose, mannose, rhamnose, and galactose) in pure culture under anaerobic conditions. Honey, which provides the host with simple carbohydrates (e.g. glucose, fructose, and sucrose) and trace amounts of secondary sugars (*e.g*. mannose, rhamnose, and galactose) (19–21). Matured pollen (beebread), provides the host with essential nutrients (i.e. amino acids, minerals, lipids), and is also rich in complex plant-derived substrates (e.g. cellobiose, xylose, and pectin)(22, 23). Therefore, the compounds selected for analysis have previously been identified, through biochemical assays, as molecules that constitute part of the honey bee diet.

When examining the individual and collective data for the utilization of glucose, fructose, and sucrose, nearly all isolates were able to utilize at least one or more of these substrate(s), with the exception of 4 taxa: *Staphylococcaceae* EBHJ3 and *Lactobacillales* MAH G07-3 within the Firmicutes and *Enterobacteriaceae* LB Tet D09 and LB Tet C11 (Figure 2, Supplemental Figure 2). At the phylum-level, at least 3 isolates within each phylum were capable of utilizing glucose, fructose, and sucrose, with varying degrees of consistency (Figure 2A, Supplemental Figure 2A). Collectively, when comparing isolates representative of taxa specific to the honey bee gut microbiota (i.e. *Gilliamella*, *Lactobacillus* Firm-5, and *Bifidobacterium*), the general trend for substrate utilization suggested that the honey bee bacteria were primed to utilize these substrates (Figure 2). Interestingly, however, we did observe differences among strains within clades in their ability to use different compounds. For example, although all *Bifidobacteria* and *Lactobacillus* Firm-4 strains used glucose, fructose, and sucrose, the *Lactobacillus* Firm-5 strain MAH H05-3 did not metabolize either monosaccharaide to a significant extent. Similarly, *Gilliamella* strains differentially processed sucrose; although both strains tested could utilize glucose and fructose, only Gamma-1 LB D06 could utilize sucrose, although to a modest extent (Figure 2).

**Figure 2.**
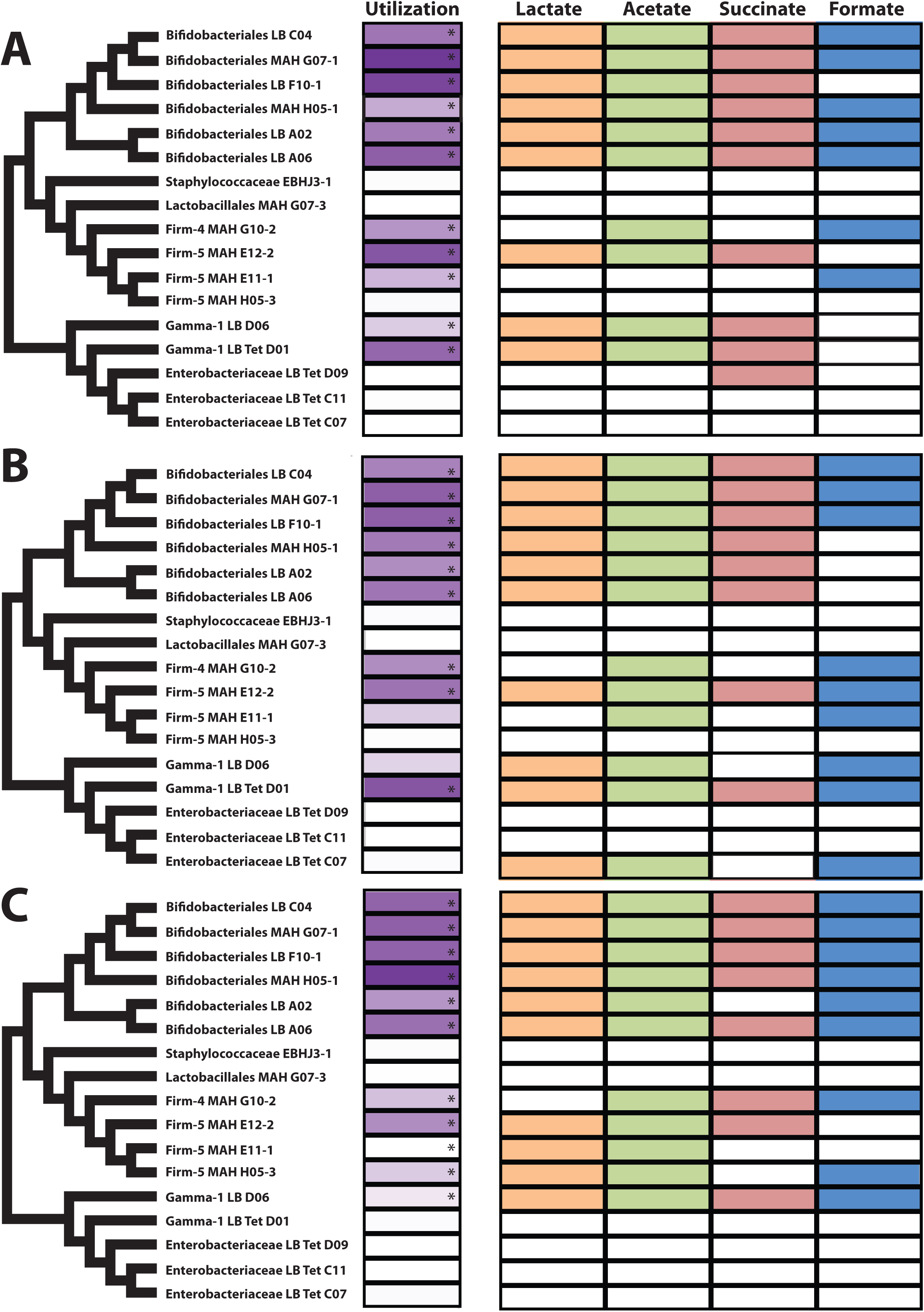
Phenotypic profile of substrate utilization by gut isolates. On the left of the figure are consensus phylogenetic trees derived from the 16S rRNA gene phylogeny. Utilization column is a heat map representation of OD590 measurements (background subtracted} from 1.4 (most purple) to zero (white). Asterisks indicate statistically significant results, compared to negative control, for glucose (A), fructose (B), or sucrose (C). Similarly, the detection of organic acids is denoted by boxes filled will the color that corresponds to a given compound detected: formate (blue), acetate (green), lactate (orange), and succinate (red), based on retention peaks on HPLC chromatograms.

### Acetate, lactate, succinate, and formate are the primary fermentation products produced by honey bee gut microbiota

To determine the soluble fermentation products produced by each individual isolate, post anaerobic metabolism, we analyzed the contents within each experimental and control well of the MT2™ plates described in the previous section using HPLC. For this analysis, we focused on the products in pure cultures with glucose, fructose, and sucrose. We chose to use simple sugars abundant in honey, rather than complex polymeric carbon sources, because complex carbon sources are degraded to simple sugars and funneled into the same catabolic pathways. With the exception of pectin, all other saccharides analyzed for utilization are mono- or di-saccharides and are only a few enzymatic reactions away from glycolysis or the pentose phosphate and Entner-Doudoroff pathways. While the ratio of fermentation products is expected to change during growth on different substrates, it is less likely that different products would be produced from different substrates. We used known standards (*e.g.*, acetate, lactate, formate, succinate) and identified peaks at the appropriate retention times within each biological sample.

HPLC analysis revealed a variety of organic acids produced by honey bee associated microbes, under anaerobic conditions. When comparing the supernatant of experimental and control wells of isolates, almost all isolates that were positive for utilizing a substrate exhibited the presence of fermentative products (as derived from retention times identical to known substrate standards, *i.e.* lactate, acetate, succinate, and formate; Figure 2). We expected to observe ethanol, acetoin, propanoate, 2,3-butanediol, based on the metabolic pathways identified in the metatranscriptomic data but did not identify other significant peaks beyond the four described above, even though the instrument is capable of detecting these compounds.

When comparing the general pattern of products produced across taxa, bacteria classified within the *Bifidobacterium* clade unanimously produced lactate and acetate, regardless of substrate. However, production of succinate and formate by this same clade was variable both across the phylogeny and across substrates (Figure 2). The Firm-5 and *Gilliamella* isolates produced these same organic acids, but with an even larger degree of variability within the clades (Figure 2). Altogether, the data for the 17 isolates supports the hypothesis that honey bee gut microbiota are primed to consume and ferment the components of the honey bee diet, but significant variation exists within strains such that no uniform profile could be ascribed to a clade or species group. For example, even if isolates of the same species used a substrate, they did not produce the same organic acids; formate was detected from some strains of *Bifidobacterium* under glucose but not under fructose, while all produced formate when given sucrose (Figure 2).

### Microbes utilize more complex carbohydrates and sugars toxic to the honey bee

We extended our analysis to plant derivatives of pollen (cellobiose, xylose, and pectin), as well as secondary sugars commonly found in honey (mannose, rhamnose, and galactose) and known to be toxic to the host in high quantities. For the plant derived carbohydrates, we observed the same trend noted for the utilization of simple sugars, with at least 2 isolates within each major taxon capable of utilizing cellobiose, xylose, and pectin. Interestingly, all honey bee specific bacteria tested could utilize pectin while more distantly related bacteria (*Staphilococcaceae, Lactobacillales, Enterobacteriaceae*) could not (Figure 3). However, we found variation in the ability of strains within honey bee associated clades in their use of other complex carbohydrates; xylose was only utilized by half of the *Bifidobacterium* strains tested and only 1 out of 4 *Lactobacillus* strains. In contrast, we observed no difference between honey bee specific and more distantly related bacteria in their ability to utilize sugars toxic to the bee (Figure 3). Again, however, we observed variation in the ability of strains to utilize these toxic sugars with only some core members able to use all of these toxic sugars and some core members unable to use any (*Bifidobacteriales* LB C04 and *Lactobacillus* Firm-4 MaH G10-2, Figure 3). Overall, these results suggesting the honey bee associated bacteria utilize mannose, rhamnose, and galactose, secondary sugars common in the honey bee diet, but toxic to the host.

**Figure 3.**
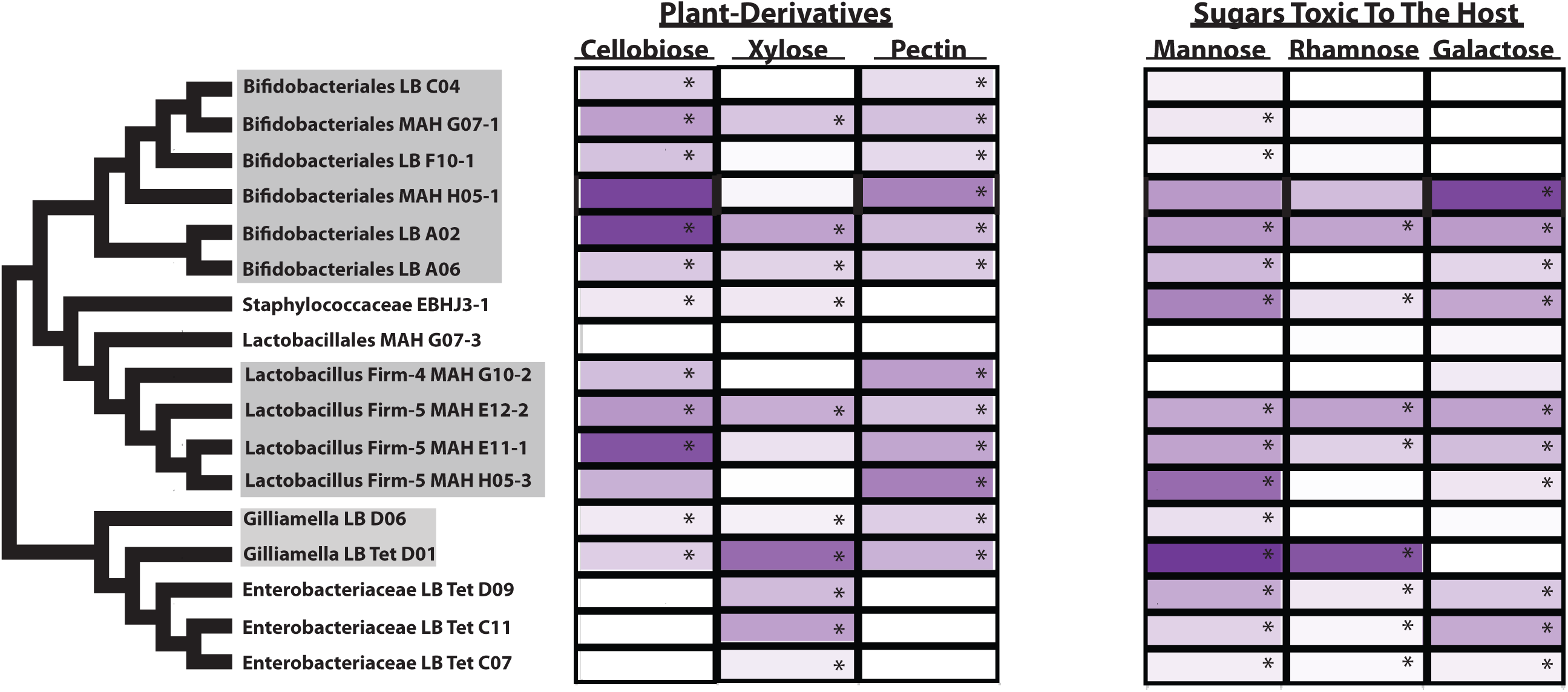
Phenotypic profile of substrate utilization by honey bee isolates. On the left are cladograms derived from the 16S rRNA gene phylogeny. Isolates highlighted in grey indicate taxa considered core to the honey bee gut, while all other isolates are representative of non-core members. The utilization of other plant-derived sugars (i.e. cellobiose, xylose, and pectin) and dietary saccharides toxic to the host (mannose, rhamnose, and galactose) is a heat map representation of OD590 measurements (background subtracted) from 1.4 (most purple) to zero (white). Asterisks indicate statistically significant results, compared to negative control.

### The honey bee gut compartments are associated with altered gene expression by the microbiota

We determined that isolates from the honey bee gut microbiota ferment sugars common to the honey bee diet into organic acids *in vitro* (Figure 2). Recent work by Zheng et al. (2017) demonstrated *in vivo*, the presence of a variety of organic acids across distinct gut chambers of the honey bee digestive tract. Based on the observations described above, we hypothesized that symbionts of the honey bee gut are likely contributing to presence of organic acids *in vivo*, and since each gut chamber houses a specific compilation of symbionts (24, 25), perhaps functional differences exist in the metabolic activity of symbionts across individual gut chambers. To address this hypothesis, we quantified the expression of genes that encode for fermentative enzymes throughout the three gut chambers of the honey bee digestive tract (i.e. crop, midgut, and hindgut (anterior to posterior)). We designed taxon-specific primers for two genes: acetate kinase (*ackA*) and L-lactate dehydrogenase (*ldh*), encoding for enzymes responsible for the production of acetate and lactate, respectively. In addition, we also designed primers for clade specific enzymes (that is, the metabolic function was only identified in one of the three phyla involved in cleaving glycosidic bonds within plant-derived compounds (*e.g.* cellulase and cellodextrinase for *Bifidobacteria*, pectate lyase for *Gilliamella*, and glucosidase for the *Lactobacillus* Firm-5 clade; Supplementary Table 1)). We used qRT-PCR on RNA extracted from each gut section to quantify the expression of these loci. For the *Gilliamella* and *Lactobacillus* Firm-5 clades, the midgut and hindgut chambers were found to be the most active gut sections (*Gilliamella* (df = 2; x^2^ = 7.05; p = 0.029), Firm-5 (df = 2; F = 3.95; p = 0.040); Figure 4A and C). In contrast, while the activity of the *Bifidobacterium* clade increases across gut sections from posterior to anterior, the trend is not significant ((df = 2; 2 = 4.36; p = 0.113); Figure 4B).

**Figure 4.**
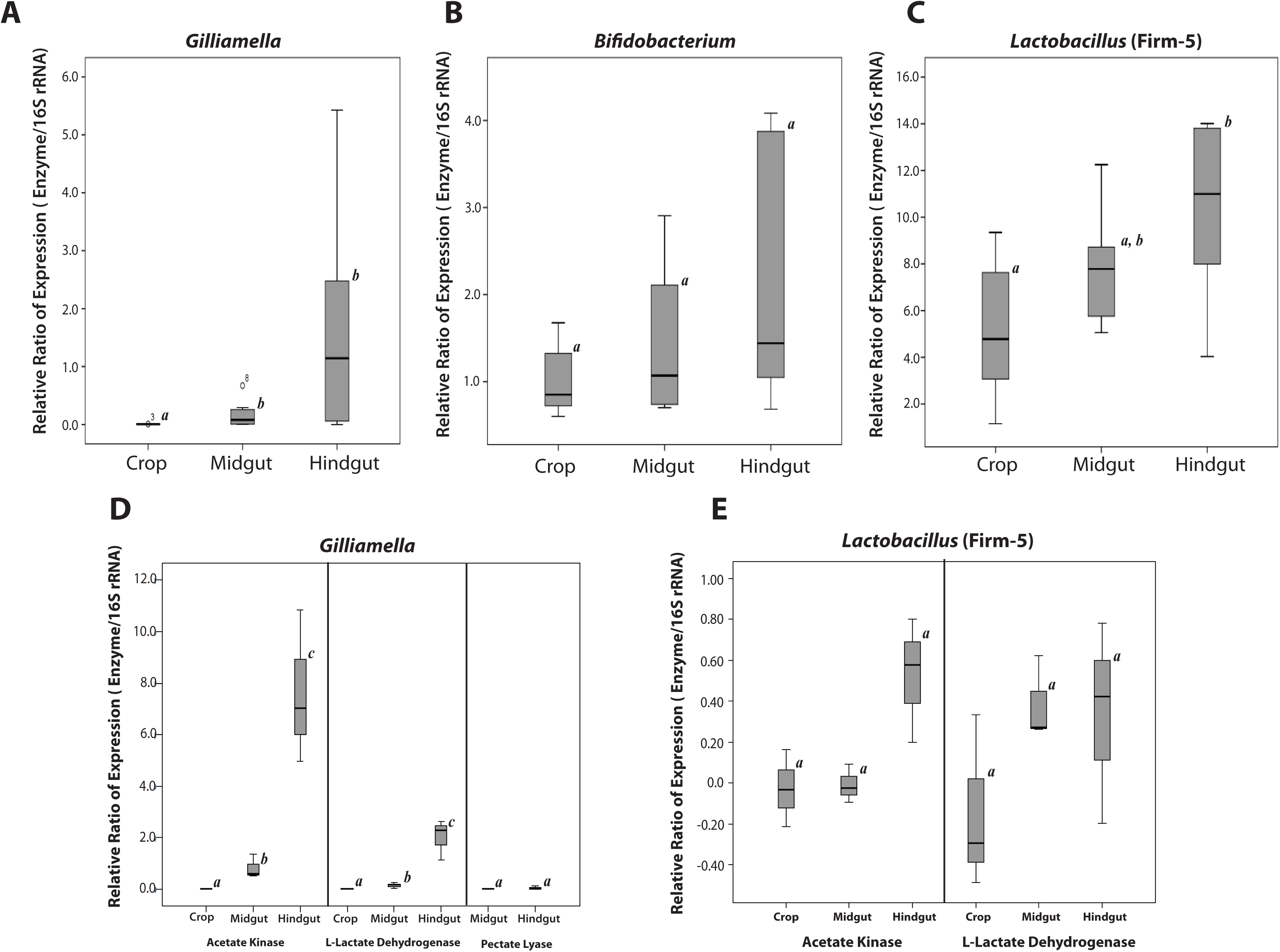
Expression of metabolic genes derived from honey bee specific taxa. Reverse transcriptase quantitative PCR (RT-qPCR) expression shown for each enzymatic genes, relative to the average 16S rRNA gene expression, across chambers of the honey bee gut. Standardized to data from the crop, the overall relative ratio of expression of enzymes for *Gilliamella* (A), *Bifidobacterium* (B), and *Lactobacillus* clades(C) were determined across gut chambers. (D-F) The relative expression data of individual genes, for *Gilliamella* (D), *Bifidobacterium* (E), and *Lactobacillus* (F), across the gut sections. For each dataset, relative expression data were compared across gut sections, using ANOVA or Kruskal-Wallis test in SPSS. For statistically significant results (p ≤ 0.05), pairwise comparisons were conducted using one-way ANOVA/post hoc or Mann-Whitney test in SPSS, significant difference (p ≤ 0.05) between gut sections are denoted with letters.

To obtain a better understanding of genes responsible for the trends seen in the *Gilliamella* and *Lactobacillus* Firm-5 clades (which both resulted in statistically significant contrasts across the digestive tract), we performed a one-way ANOVA/Tukey-HSD to analyze the differences in variation of relative gene expression across gut sections. For *Gilliamella*, the analysis of ackA and ldh expression indicated that expression levels significantly increased across gut sections in the following order: crop<midgut<hindgut (Figure 4D). Pectate lyase was detected at equal levels of expression in the midgut and hindgut, but no amplification was detected in the crop (Figure 4D). In contrast to *Gilliamella*, we saw no pattern to the expression of ldh or ackA by *Lactobacillus* Firm-5 across the gut sections (Figure 4E). Importantly, we were not able to detect β-glucosidase and cellulase gene expression from Firm-5 and *Bifidobacterium* clades, in any gut section, suggesting that they are not active in the honey bee gut.

## Discussion

The honey bee gut microbiota is a deceptively simple community – although dominated by 8-10 species (26) and transcriptionally dominated by only 3 phyla (5), the strain diversity found within the gut is high (16). Here we show that strains within three transcriptionally dominant clades associated with the bee are functionally distinct and that there is indeed, no strict metabolic rule that describes each clade. Although overall, the microbes seem primed to utilize carbohydrates in the honey bee diet, expressing enzymes capable of cleaving glycosidic bonds and fermenting these substrates, strains within each clade of core microbes investigated here differ in their ability to use different simple sugars, sugars toxic to the bee, or complex carbohydrates found in the bee diet (Figures 2, 3). They also differ in their production of organic acids, the production of which are suspected to contribute to honey bee metabolism. The results presented in this study reveal that bacterial taxa associated with the honey bee gut are differentially present and metabolically active throughout the three major gut chambers, with particular enrichment in the hindgut. Of the three taxa assayed for gene expression, the *Gilliamella* clade of bacteria was the most active, showing exceptional expression of its acetate kinase gene in the mid- and hindgut, a result supported by previous metatranscriptomic analyses.

The utilization results presented in this study are supported by previous observations of gross pH changes when honey bee isolates are cultured (9, 11–14) and previous measures of acetic and lactic acid production from a few pure cultures (27, 28). Although our results are well supported by predicted models based on the genomic evidence, we did not detect the production of specific fermentative waste products predicted (e.g. ethanol, acetoin, propanoate, 2,3-butanediol)(5). The lack of detection of these products in our analysis could be due to the following: 1) a complete lack of production under the *in vitro* conditions provided 2) or production below the detectable range of the UV and refractive index detectors used.

In addition to analyzing the utilization of the substrates mentioned above, we also examined the ability of isolates to utilize pectin, which is a heteropolysaccharide, commonly found in the primary cell wall of terrestrial plants, such as pollen grains. In our analysis, we identified multiple isolates within the *Gilliamella, Lactobacillus* Firm-5, and *Bifidobacterium* clades able to utilize pectin (Figure 3). Importantly all tested core microbiome strains utilized pectin in our assays – it was the only consistently utilized carbohydrate tested – and more distantly related organisms (e.g. *Staphylococcaceae, Lactobacillales, Enterobacteriaceae*) were unable to use it. Previous work has shown that *Gilliamella* isolates are capable of degrading pectin, possessing a pectate lyase gene capable of cleaving de-esterified pectin (7). In addition to *Gilliamella*, we also report utilization of pectin and/or its derivatives by isolates within the Firm-4/-5 and *Bifidobacterium* clades, observations supported by a recent publication that observed growth of several core gut isolates in minimal media supplemented with pollen extract (15). These results suggest that multiple constitutes of the honey bee microbiota may be capable breaking-down and utilizing plant material abundant in the host’s diet, an idea that has previously been hypothesized (10). Indeed, the genome of *Bifidobacterium asteroides* has an annotations for glycosidases and a pectinesterase, enzymes capable of degrading pollen (6). Contrary to *Bifidobacterium*, genomes from bacteria in the *Lactobacillus* Firm-5 clade currently have no annotated pectinases or glycosidases (8), but these and many of the genomes of honey bee specific bacteria possess a plethora of hypothetical genes of unknown function that might aid in the degradation and metabolism of pectin (9). Importantly, although the pectin used in our *in vitro* assays was of high quality, it is not a completely uniform nor pure substance and therefore it is possible that trace amounts of other compounds might have been present and metabolized by the honey bee specific microbes. Further experiments need to be conducted to confirm the pectinase activity of the isolates analyzed. It should also be noted, that the conditions provided in the utilization assay likely do not support the growth of all 17 isolates as the salt solution lacks several elements required for growth. However, although the assay does not rely on growth, it is possible that the utilization patterns observed for individual isolates could potentially alter under alternative conditions conducive for growth.

Recent important work has identified a link between the presence of canonical gut symbionts and weight gain in honey bees; organic acids, which are presumed to be bacterially derived, are thought to contribute to the observed influence symbionts to host health (28). The fact that these microbes live within the bee gut and are consistently associated with honey bees has led to the assumption that microbes isolated from the bee are mutualists in this context – that they benefit the bee in some way. However, supplementation of the bee diet with one of these core members (*Snodgrasella)* results in an increased susceptibility to parasites (29) and another (*Frischella*) results in scab formation in the intestine (7). The converse can also be true; non-core members of the bee can provide a benefit as well. *Lactobacillus kunkeii*, an environmentally acquired microbe, protects from pathogens (30) and although *Gilliamella* strains have been found to use sugars that are toxic to the bee (31), here we show that both non-core bacteria and core microbes can utilize these sugars.

The difference in the enzymatic activity pattern between members of the honey bee microbiota suggests they might occupy different niches in the honey bee food, digestive tract, or built environment. The idea that niche partitioning might exist in the gut of the honey bee is physically plausible, as the alimentary system in the honey bee is subdivided into three distinct chambers. Indeed, our analysis of gene expression (qRT-PCR) suggests that expression of different enzymes can differ across gut sections, genes, and taxa. It should be noted, however, that we could not detect the expression of genes responsible for the breakdown of complex plant-derived molecules (i.e. glucosidase, cellodextrinase, and cellulase). These results indicate that these genes are either completely inactive, or are expressed at levels below detection. In either scenario, the data suggest that perhaps complex carbohydrates are either present at minimal levels in honey bee gut, or that the taxa analyzed in this study thrive on other resources abundant in the environment (i.e. glucose, fructose, sucrose, and/or other plant derivatives) by expressing other enzymatic loci capable of contributing to catabolism.

Our work suggests a dominant profile for *Gillamella in vivo.* Of the three analyzed taxa, the *Gilliamella* clade was the most active in our analysis, with the *Bifidobacterium* and *Lactobacillus* Firm-5 clades being roughly equal; this result fits well with previously published transcriptomic data, where the gamma-proteobacteria dominated the transcriptional profile of the gut (5). When assaying the relative expression of certain genes (i.e. ackA and ldh) for the *Gilliamella* clade (Figure 4A, D), we saw highest levels of expression in the hindgut, suggesting that *Gilliamella* may function differently in each gut chamber or may be particularly active in the hindgut. Additionally, within each gut section, *Gilliamella* consistently expresses more ackA than ldh, which might suggest a preference for this taxon to produce acetate, a byproduct produced *in vitro* by *Gilliamella* isolates under nearly all tested conditions (Figure 2). It is intriguing that *Gilliamella* dominates transcriptomic analyses of the honey bee and related species are found in bumble bees (8). *Gilliamella* and other proteobacterial species in the honey bee require direct social contact between bees beyond mere trophallaxis (i.e. fecal exposure) to achieve transmission (32). The fact that the honey bee workers acquire this important symbiont through their social interactions, and that *Gilliamella* dominates the functional profile of the honey bee worker gut suggests it to be an important, if not critical, part of the honey bee core microbiota.

Recent work elegantly demonstrated through metabolomics that honey bees primarily colonized by individual bacterial strains of taxa core to the gut microbiota show depletion in plant derived substrates and an enrichment in fermentation product present in the digestive system, compared to bees with a depleted gut microbiota (33). These results are promising, as they demonstrate substrate utilization by individual core taxa and enrichment of metabolites within the gut *in vivo.* However, our work here suggests that the choice of strain used in colonization experiments may substantially influence results; not all *Gilliamella apicola* strains produce the same organic acids under the same conditions. It is perhaps the case that the diversity of strains found within the honey bee gut complement each other, providing a range of organic acids under most conditions and diets. This hypothesis awaits testing. Nevertheless, it is obvious that individual taxa found in the core microbiome of bees might not accurately reflect the metabolic dynamics occurring in the gut of a honey bee with a natural microbiota, where microbe-microbe and microbe-host interactions likely govern niche partitioning and microbiome function.

## Acknowledgements

We thank anonymous reviewers for their feedback on early stages of this manuscript.

